# Enhancing STED Microscopy via Fluorescence Lifetime Unmixing and Filtering in Two-Species SPLIT-STED

**DOI:** 10.1101/2025.07.17.661952

**Authors:** Andréanne Deschênes, Antoine Ollier, Marie Lafontaine, Albert Michaud-Gagnon, Jeffrey-Gabriel Steavan Santiague, Anthony Bilodeau, Christian Gagné, Paul De Koninck, Flavie Lavoie-Cardinal

**Affiliations:** CERVO Brain Research Centre,Université Laval; Institute for Intelligence and Data, Université Laval; Department of Electrical and Computer Engineering, Université Laval; Department of Biochemistry, Microbiology and Bio-informatics, Université Laval; Department of Psychiatry and Neuroscience, Université Laval

## Abstract

Simultaneous super-resolution imaging of multiple fluorophores remains a major challenge in STimulated Emission Depletion (STED) microscopy due to spectral overlap of STED-compatible fluorophores. The combination of STED microscopy and Fluorescence Lifetime Imaging Microscopy (FLIM) offers a powerful alternative for super-resolved, multiplexed imaging of biological samples but is hindered by lifetime convergence at high depletion powers. Here, we present an analysis method, two-species Separation of Photons by LIfetime Tuning (SPLIT)-STED, that uses a linear system of equations in phasor-based STED-FLIM to enhance both fluorophore unmixing and spatial resolution. It defines the fluorescence signal as a mixture of three lifetime components: the two target fluorophores and a short-lifetime contribution from undepleted peripheral fluorescence photons. Two-species SPLIT-STED disentangles overlapping lifetimes and selectively filters low-resolution signal. The method enables accurate unmixing of spectrally overlapping fluorophores and, by enhancing resolution through lifetime-based filtering, allows the use of lower depletion powers, thereby improving fluorescence lifetime separation.

## 1 Introduction

Multiplexed super-resolution microscopy is essential for characterizing molecular interactions and structural remodeling within subcellular compartments. In multi-color STED microscopy, multiplexing is typically achieved through spectral separation of fluorescent labels [1]–[5], a strategy that is often limited by spectral overlap and co-excitation of fluorophores [1], [6]. As an alternative, combining STED with fluorescence lifetime imaging (FLIM) enables fluorophore discrimination based on lifetime distinction [7]–[14].

Depletion of the fluorescence signal in STED microscopy modifies the measured lifetime distribution by creating a multi-exponential fluorescence decay. It results from the competition between the fluorescence and the stimulated emission depletion processes. This effect is more pronounced at the periphery of the Point Spread Function (PSF) and thus of the nanostructures in STED images [9], [15]–[17]. Resolution improvement based on lifetime filtering of STED microscopy images is known as SPLIT-STED and was first introduced and demonstrated with Continuous Wave (CW)-STED-FLIM by Lanzan’o *et al*. [17]. SPLIT-STED was then adapted to different experimental paradigms: CW-STED-FLIM with two-photon excitation [18], CW-STED-Fluorescence Correlation Spectroscopy (FCS) [19], and pulsed STED-FLIM measured in the frequency [20] and the time [21], [22] domains. Recently, SPLIT-STED was also combined with machine learning to make it usable at lower photon counts [23] and generalized to be used in image-scanning microscopy [24]. A similar method has been implemented by Leica Microsystems as Tau-STED [25], which is being used for different biological assays [7], [26]–[31]. Until now, these resolution improvement methods have been used for single-species STED-FLIM.

Analytical methods have been developed to extract the lifetime of fluorescent species contributing to the STED-FLIM signal. Exponential model curve fitting has been employed for various applications in STED-FLIM, such as separating the contribution of fluorescent species [9], [11], evaluating FRET efficiency [32], and improving spatial resolution and signal to noise ratio (SNR) [21], [33]. The performance of curve fitting approaches heavily depends on the number of measured fluorescence photons in each pixel [9], [11] becoming increasingly complex for higher numbers of fluorescent species [34]. The phasor approach, an alternative analysis method for FLIM analysis, was introduced by Digman *et al*. [35] providing a graphical global view of the distribution of fluorescence decays at each pixel. More precisely, the fluorescence lifetime of a single pixel can be represented as a point in the phasor space. In STED-FLIM, binary selections [8], [20] and linear equation systems [7], [12]–[14], [30] have been used to discriminate between up to three fluorescent species using phasor plot analysis. Although increasing STED depletion power improves spatial resolution [36], it reduces the lifetime difference between fluorophores, thereby complicating their separation in STED-FLIM [7], [8], [14], [30].

We propose a phasor-based analysis approach for two-species SPLIT-STED microscopy that leverages a linear system of equations to enhance both fluorophore unmixing and spatial resolution in STED-FLIM. Our method assigns detected photons to three distinct components: the two target fluorescence lifetimes and a third component representing short-lifetime undepleted fluorescence photons at the periphery of the structures of interest.

## 2 Materials and Methods

### 2.1 Dataset acquisition

#### 2.1.1 Hippocampal rat neuronal cultures

Rat neuronal hippocampal cultures were prepared in accordance with procedures approved by the animal care committee of Université Laval. We used a previously established protocol, optimized for low density neuronal cultures on glass coverslips [37]. Hippocampi were dissected from postnatal rats (P0-P1) and subjected to both enzymatic digestion with papain (12 U/mL; Worthington Biochemical Corporation) and mechanical dissociation via trituration. Following dissociation, cells were plated onto PDL-Laminin coated glass coverslips (12 mm diameter) in a 24-well plate [38]. Fetal bovine serum (2%; Hyclone) was added to the culture medium at the time of plating. The PDL-Laminin coating allowed to maintain healthy cultures for at least 3 weeks at a neuronal cell density of 200 cells/mm^2^, enabling the isolation of axonal and dendritic projections for super-resolution microscopy experiments [37].

The cultures were maintained in Neurobasal medium (Thermofisher) supplemented with B-27 (50:1), penicillin (50 U/mL), streptomycin, (50 µg*/*mL) and L-GlutaMAX (0.5 mM; Thermofisher). After five days in culture, half of the media was replaced with fresh media devoid of serum and containing Ara-C (5 µM; Sigma-Aldrich) to inhibit non-neuronal cell proliferation. Subsequently, the cultures were fed twice a week by replacing half of the growth medium with serum- and Ara-C-free medium.

#### 2.1.2 Fixation and immunostaining

Cultured neurons were fixed at 14 to 21 days in vitro (DIV) using a freshly prepared solution consisting of 4% paraformaldehyde (PFA) supplemented with 4% sucrose, 100 mM phosphate buffer, and 2 mM Na-EGTA. The fixation solution was maintained at 37°C. Optimized fixation times of 10 minutes were used for synaptic proteins (PSD95, Bassoon, Homer1) and 20 minutes for cytoskeletal proteins (*α*-Tubulin, F-Actin, *β*II-Spectrin) [37], [39]. After fixation, the cells were washed three times for 5 minutes with Phosphate-Buffered Saline (PBS) containing 100 mM glycine. Prior to immunostaining, the cells were permeabilized in a normal goat serum (NGS) (2%; Cedarlane, CL1200-100) blocking solution of PBS supplemented with Triton X-100 (0.1%; Fisher Scientific, BP151-100). Incubation with primary antibodies (PAB) and secondary antibodies (SAB) was carried out in the NGS blocking solution (see Table S1 for antibody details). PABs were incubated for 2 hours at room temperature (RT), followed by three washes in PBS. The SABs were incubated for 1 hour at RT and then washed three times with PBS. Finally, coverslips were mounted on microscopy glass slides using polyvinyl alcohol mounting medium with DABCO (Sigma-Aldrich, 10981).

#### 2.1.3 STED-FLIM microscopy

STED-FLIM images were acquired using an Expert Line STED microscope (Abberior Instruments GmbH) equipped with a 775 nm depletion laser (40 MHz, 1.2 ns pulse duration; MPB Communications), four Avalanche Photodiode Detectors (APD) (Excelitas, SPCM-AQRH-13) and four pulsed excitation lasers (485 nm, 518 nm, 561 nm, and 640 nm). A Time Tagger Ultra 8 (Swabian Instruments GmbH) was connected to two APDs to enable Time Correlated Single Photon Counting (TCSPC) measurements. Control of the Time Tagger was performed using the Imspector software (version: 16.3.15513, Abberior Instruments GmbH) of the STED microscope. STED-FLIM was performed using red and far-red emitting dyes, excited at 561 and 640 nm respectively, and depleted with a single depletion laser beam at 775 nm. The time-resolved fluorescence signal was detected on the APDs using FF02-615/20-25 and ET685/70 (red; Semrock, far-red; Chroma) filters. The excitation laser pulse synchronization signal and the electrical output of the APD detectors were connected to the Time Tagger to assign time tags to the excitation laser pulses and fluorescence photon counts. The delay between the excitation laser pulses and the arrival time of the photons on the APD were computed and compiled into fluorescence decay histograms. Additionally, line and frame triggers generated by the microscope were connected to the Time Tagger to assign the tagged photons to the correct pixel’s histogram. The Olympus autofocus unit (Olympus, IX3-ZDC2-830) was used to stabilize the sample focus during the image acquisition process. Excitation laser powers and line repetitions were set to maximize photon counts per pixel, while minimizing pile-up effect [40]. The histogram settings were 250 bins over 20 ns (80 ps per bin). For the STED-FLIM experiments the pinhole was set to be (*∼*1 Airy unit at 100X magnification for red fluorescence). The imaging parameters, including laser powers measured at the objective’s Back Focal Plane (BFP), are presented in Table S2.

#### 2.1.4 Imaging automation

A Python script using the SpecPy [41] and Abberior-STED [42] packages was created to automate image acquisition and the selection of the regions of interest (ROI) in the Imspector software (version: 16.3.15513, Abberior Instruments GmbH). Automated acquisition was used to set the depletion power from a list of randomly shuffled depletion powers and acquire predefined sequences of images for each ROI. Automated acquisition scripts can be found here: https://github.com/FLClab/2-Species-SPLIT-STED.

### 2.2 Histogram fitting

Histogram fitting was performed with the Maximum Likelihood Estimation (MLE) method which compares FLIM histograms to a predefined mono-exponential function model (equation 1) [43]. By adjusting the parameter *τ* (equation 1), it seeks to minimize the difference between the model and the observed data. The optimization process relies on the `SciPy`library’s [44] `minimize`function, using the `Sequential Least Squares Programming (SLSQP)`algorithm [45]. MLE fitting requires an initial estimate of *τ* and established boundaries, which were set between 0.1 and 5 ns. The output of the MLE fit is the estimated fluorescence lifetime, *τ*, and when applied to multi-exponential data, MLE estimates the mean lifetime [46].

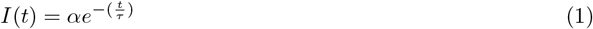

Considering that the measured fluorescence lifetimes (1-5 ns) were much longer than the Full Width at Half Maximum (FWHM) of the Instrument Response function (IRF) (634 ps) of our system, we performed tail-fitting of the decay histogram [47]. Thus, before applying histogram fitting, the 20 first temporal bins were removed. The resulting histograms were normalized by dividing the photon counts per bin by the total number of photons. Histogram fitting was performed on each pixel which contained more than five photons.

Pixels containing less than five photons were assigned a lifetime of 0 ns [9]. To display the pixel lifetime values on the intensity images, a color-code was assigned to the measured lifetime and the pixel’s intensity was used to determine brightness. The mean fluorescence lifetime of each image is calculated using a summed histogram of all photons from pixels with more than five detected photons.

### 2.3 Phasor distribution calculation

The phasor analysis method translates photon arrival times into a two-dimensional distribution of points in a polar coordinate system, referred to as a phasor plot, using a sine-cosine transform [35]. Each pixel in the FLIM image corresponds to a unique point on the phasor plot.

The pixel coordinates in the phasor space (*g, s*) correspond to the sine-cosine transform of the fluorescence decay histogram (*I*_*i,j*_(*t*)). The transforms are calculated using [35]:

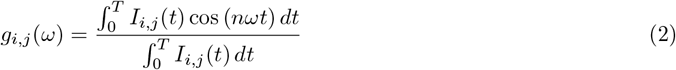

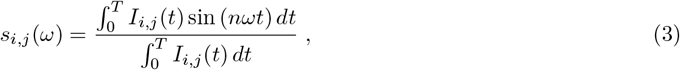

with *ω* corresponding to the angular laser pulse repetition frequency (*ω* = 2*π \**40Mrad/s for a 40 MHz laser), *i, j* to the X and Y coordinates of a pixel of the original microscopy image, *T* to the period between the laser pulses, and *n* to the harmonic frequency (set to one in this method).

Fluorescence decay histograms are convolved by the IRF of the imaging system. The result of this convolution in phasor space is a shift of the position of the phasor distribution in the universal semicircle [48]. To remove this shift and return the phasor distributions to the universal semicircle, the phasor coordinates were calibrated by applying a displacement in polar coordinates (*m,ϕ*)(Figure S2B, equations 4 and 5) [48]–[50]. The size of this displacement was determined using the experimental IRF, measured with the backscattering of 150 nm gold nanospheres [51]. The measured gaussian IRF has a FWHM of 634 ps (Figure S2A).

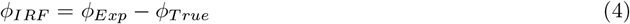

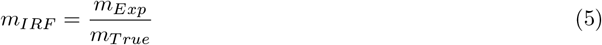

where *ϕ*_*IRF*_ is the phase shift and *m*_*IRF*_ is the radial modulation factor. The size of these displacements (*ϕ*_*Exp*_ and *m*_*Exp*_) are determined using the centroid of the phasor distribution of the IRF measurement.

The calibrated phasor coordinates were then filtered either using two-dimensional convolution by a median filter as in [22] (kernel size: 3, smooth factor: 0.2) or the Complex Wavelet Filter (CWF) filter [52] (50 neighbors and 2 levels). The effect of the two filtering techniques on a phasor distribution and the resulting SPLIT-STED images are shown in Figure S4.

Fluorescence lifetimes that can be described by a single exponential decay are located in phasor space on the universal semicircle (circle with a radius of 0.5 centered at *g* = 0.5, *s* = 0). Measured lifetimes best described by multi-exponential decays are distributed within the semicircle and along a linear trajectory joining their mono-exponential components. As a result of the principle of linear combination of phasors, the relative contributions of the mono-exponential components can be quantified based on the position along the connecting line.

### 2.4 Single-species SPLIT-STED

The algorithm described by Tortarolo *et al*. [22] was reproduced to perform single-species SPLIT-STED. Each image was separated into two lifetime fractions: 1) a longer lifetime in the central region of the imaged structure, and 2) a shorter lifetime in its periphery [17].

The phasors were filtered using the CWF approach [52] to reduce the spread of the phasors of the Confocal-FLIM and STED-FLIM images while preserving high spatial frequency structures (Figure S4). The contributions from the short and long lifetimes were separated using a linear decomposition approach for each of the STED-FLIM image’s pixels [22]. For single-species SPLIT-STED we measured the centroid position of the confocal-FLIM phasor distribution and defined *P*_*n*_ as its nearest point on the universal semicircle. Each point of the STED-**FLIM** image’s phasor distribution was orthogonally-projected onto the line connecting *P*_*n*_ and the *g*=1, *s*=0 limiting point, *P*_*l*_ (Figure 1C). The coordinate of each projected point was converted to a parametric coordinate on the SPLIT-STED trajectory line. This parametric coordinate scaled between zero for a pixel having only the short lifetime (positioned at *P*_*l*_) and one for a pixel having only the long lifetime (positioned at *P*_*n*_)(Figure 1C). The parametric coordinates were rescaled to match the spread of the STED-FLIM image’s phasor distribution. Two points on the linear trajectory were used as *P*_1_ and *P*_2_ between which the parametric coordinates were normalized to be between zero and one. To determine the position of *P*_1_ and *P*_2_, two centroids were calculated for the STED image’s elongated phasor and were orthogonally projected onto the linear trajectory. Figure 1D illustrates the single-species SPLIT-STED process.

**Figure 1.**
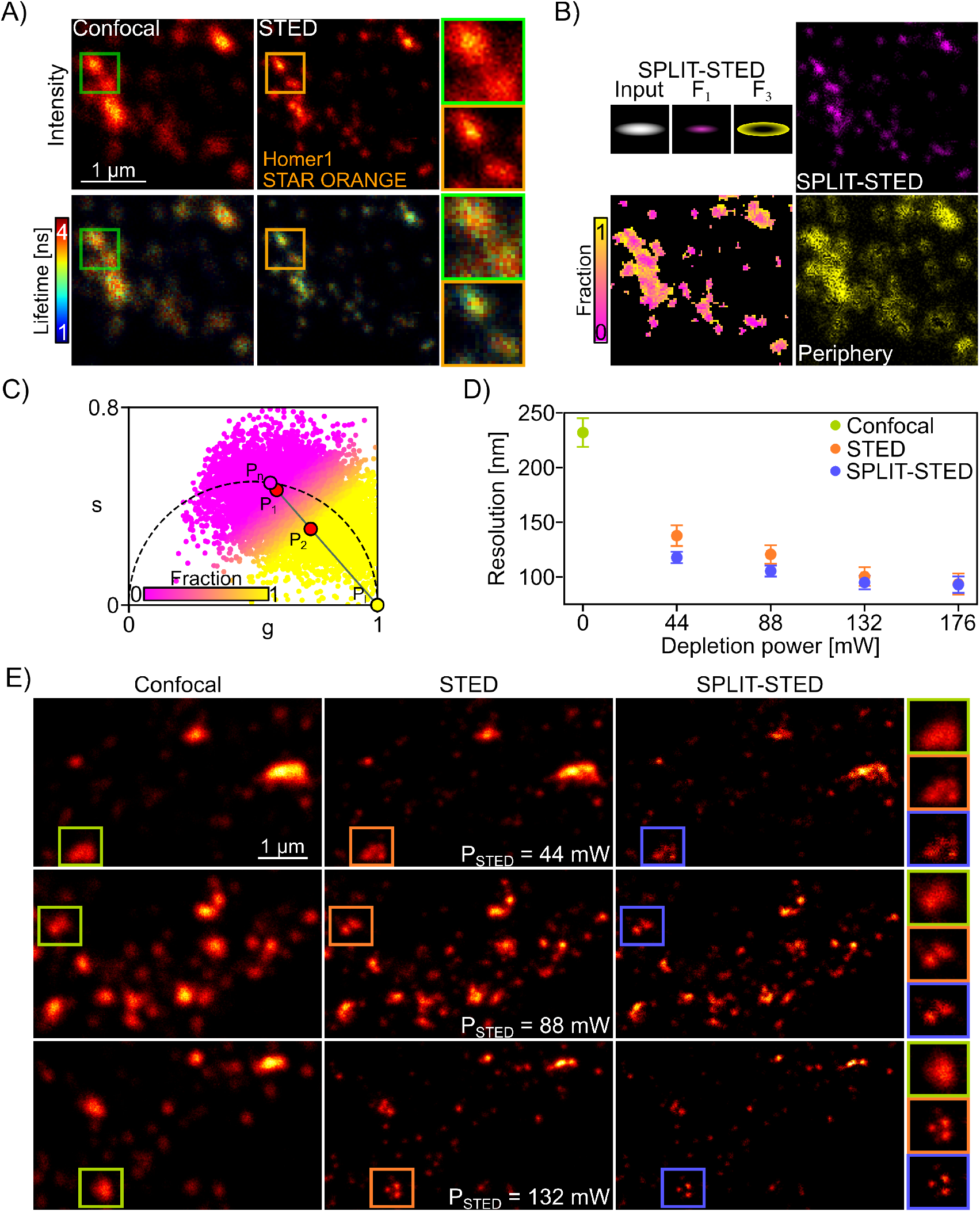
Single-species STED-FLIM and SPLIT-STED. (A) Confocal and STED images (*P*_*ST ED*_=88mW) of the synaptic protein Homer1 labelled with STAR ORANGE. The images are color-coded for the photon counts (top) and the measured fluorescence lifetime (bottom). (B) SPLIT-STED separates the shorter lifetime component at the border of the structures (*F*_3_, yellow) from the longer component at the center (*F*_1_, magenta). The resulting SPLIT-STED images of the *F*_1_ fraction is characterized by an enhanced spatial resolution. (C) Color-coded phasor plot of the lifetime distribution from the STED-FLIM image shown in (A). The trajectory line between the confocal lifetime (*P*_*n*_) of STAR ORANGE and limiting point *P*_*l*_ is used to assign pixels to their corresponding fraction (Material and Methods). The points in the phasor plot are color-coded based on their position on the trajectory line relative to the two main lifetime components (Fraction 1: *P*_1_, Fraction 3: *P*_2_). (D) Mean resolutions (*±* standard Deviation (STD)) measured using the decorrelation method on images of Homer1-STAR ORANGE. SPLIT-STED improves the spatial resolution for a given depletion laser power (Number of images : 6 - 44 mW; 9 - 88 mW; 9 - 132 mW and 10 - 176 mW). (E) Representative confocal, STED and resulting SPLIT-STED images of Homer1 acquired with different depletion powers.

### 2.6 Phasor distribution characterization

The spread of the distribution in a phasor plot inversely scales with the number of photon counts in each pixel and with the heterogeneity of lifetimes in the image [35], [52]. To investigate the impact of the depletion power on the phasor distributions of STED-FLIM images, the position and spread of phasor distributions were characterized. To do so, an ellipse was fit to the 70^th^ percentile of the phasor distribution. The shortest distance between the ellipses of phasor plots obtained for different fluorophores and at different depletion powers were computed to characterize the difference between phasor distributions. The distance measurement was used as an indicator of the difficulty to discriminate between two fluorophores in the phasor space (Figure S5B).

### 2.6 Two-species Confocal- and STED-FLIM

To separate the contributions of 2 lifetime components in STED-FLIM images, linear separation in phasor space was performed [7], [12], [13], [30]. We automated the position calculation of the two pure species in phasor space to remove user bias.

We used the centroids of the phasor distributions of single-staining sample images as reference points (*P*_1_ and *P*_2_) for the pure species. Each point of the phasor distribution of the two-species STED-FLIM image was orthogonally projected onto the line connecting the reference points (Figure S6). The coordinate of each projected point was converted to a parametric coordinate, *c*, on the line (*c* = 0 at *P*_1_ and *c* = 1 at *P*_2_). The parametric coordinates’ values were clipped to be between zero and one. These parametric coordinates represent the fractional composition of one species (*f*_1_). The fractional composition of the other species is obtained using *f*_2_ = 1 − *f*_1_. The intensity of the two-species image was then multiplied by these fractional compositions to obtain unmixed images (*F*_1_,*F*_2_).

### 2.7 Two-species SPLIT-STED

The general principle of two-species SPLIT-STED is to separate the signal in a STED-FLIM image into 3 fractional components using a reference triangle in phasor space (fluorophore 1 - *f*_1_, fluorophore 2 - *f*_2_ and undepleted photons - *f*_3_). The centroids of the phasor distributions from single-species Confocal-FLIM images are used to determine the reference position of each fluorophore (*P*_1_ and *P*_2_) on the universal semicircle. Three STED-FLIM images at varying depletion powers (44, 88, and 132 mW for the red channel; 22, 44, and 66 mW for the far-red channel) were acquired and a linear fit was performed to connect the centroid positions. The intersection of the resulting linear trajectories defines a point (*P*_4_), and the third reference point (*P*_3_) of the triangle is assigned to the closest location to *P*_4_ within the universal semicircle. The reference triangle for two-species SPLIT-STED is built using the *P*_1_, *P*_2_ and *P*_3_ reference points (Figure S7). The coordinates of *P*_1_, *P*_2_, and *P*_3_ were used to build the system of 3 equations described in [53]:

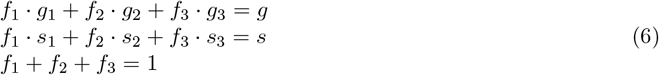

Where *g*_1,2,3_ and *s*_1,2,3_ are the coordinates of the triangle’s vertices (*P*_1,2,3_), while *g,s* are the coordinates of a pixel in phasor space for which the fractional composition (*f*_1_, *f*_2_, *f*_3_) is being evaluated.

The system of equations was solved for each point in the phasor distribution of the two-species image. Each point (pixel) was thus associated with a proportion of the three fractions (*f*_1_,*f*_2_,*f*_3_) based on its position in phasor space in relation to *P*_1_,*P*_2_ and *P*_3_. The value of the third fractional component *f*_3_ was rescaled to be between zero and one between the two centroids of the two-species image phasor distribution. The first 2 fractional components (*f*_1_,*f*_2_) were then adjusted proportionally to maintain complementarity (*f*_1_ + *f*_2_ + *f*_3_ = 1). The intensity of each pixel in the STED-FLIM image was multiplied by its corresponding fractional components (*f*_1_,*f*_2_,*f*_3_) to build the fraction images (*F*_1_,*F*_2_,*F*_3_). The photons associated with *F*_3_ (undepleted photons) are filtered while the photons associated with *F*_1_ and *F*_2_ are used to generate the unmixed images for each fluorophore.

### 2.8 Performance evaluation on a synthetic dataset

A synthetic dataset of two-species STED-FLIM images was created by combining single-species STED-FLIM images that were acquired with the same imaging parameters [9]. Each single-species STED-FLIM image was used as a ground truth of the spatial distribution of its species. This provided FLIM images of two fluorophores with a corresponding ground truth that could be used to quantitatively assess the unmixing performance of different analytical approaches. The synthetic images were built by summing photon counts pixel- and bin-wise. Zeros were added to pad regions of image size mismatch.

The NanoJ-Super-resolution QUantitative Image Rating and Reporting of Error Locations (SQUIRREL) [54] metric was used to measure the error between the ground truth STED image and the unmixed fraction image. NanoJ-SQUIRREL error maps were produced by calculating the pixel-wise Squared Error (SE) between the convolved images. The spatial resolution of the ground truth STED images and separated fraction images were calculated using the decorrelation method [55].

### 2.9 Statistical Analysis

Normality of the data was assessed with the Shapiro-Wilk test. For normally distributed data, a one-way ANOVA was used to test the null hypothesis that there were no differences among group means. When the ANOVA indicated significance, post hoc pairwise comparisons were performed using the t-test. For non-normally distributed data, the Kruskal–Wallis H test was used to assess the null hypothesis that all group medians are equal. If the null hypothesis was rejected, post hoc pairwise comparisons were conducted using Dunn’s test. Statistical tests were performed using the SciPy [44] and scikit-posthocs [56] libraries in Python.

## 3 Results

### 3.1 Single-species SPLIT-STED

We first characterized the effect of the depletion process in STED microscopy on the lifetime distribution of STED-FLIM images (Figure 1A, Figure S3). In comparison to Confocal-FLIM images, the fluorescence lifetime distribution of STAR ORANGE is modified in STED-FLIM images [9], [14], [32]. More specifically, a lifetime reduction is observed at the borders of the synaptic protein nanoclusters in the STED-FLIM images, which is used for resolution enhancement in the SPLIT-STED approach [16], [21](Figure 1A,B Figure S3). We applied the SPLIT-STED strategy on STED-FLIM images of Homer1-STAR ORANGE to separate the measured FLIM signal into two fractions that are associated with a shorter lifetime component (border) and a central longer lifetime component (Figure 1B,C). As demonstrated in previous studies, this approach increases the effective spatial resolution in comparison to conventional STED imaging, with stronger effects observed at lower depletion powers (Figure 1D). Thus, SPLIT-STED enables the reduction of the depletion power needed to resolve synaptic nanodomains of Homer1 (Figure 1E).

### 3.2 Two-species SPLIT-STED

We next evaluated the application of SPLIT-STED to two-species STED-FLIM imaging. Lifetime unmixing of two fluorophores in STED-FLIM images is increasingly challenging when raising the depletion power as it reduces the mean lifetime difference between a given fluorophore pair (Figure 2A-B, Figure S5). This is related to the mean lifetime reduction associated with the stimulated emission process [15], [17]. We observed that an increase in the depletion laser power results in an improvement in the spatial resolution. However, it is also associated with an important reduction (18%) of the mean lifetime difference between two fluorophores (STAR ORANGE and CF594, Figure 2C). In the phasor space, we observed an increase in the ellipticity and a shift to the center of the universal semicircle for the lifetime distribution of STED-FLIM images (Figure S1). It results in highly overlapping lifetime distributions for two-species STED-FLIM (Figure 2D, Figure S5).

**Figure 2.**
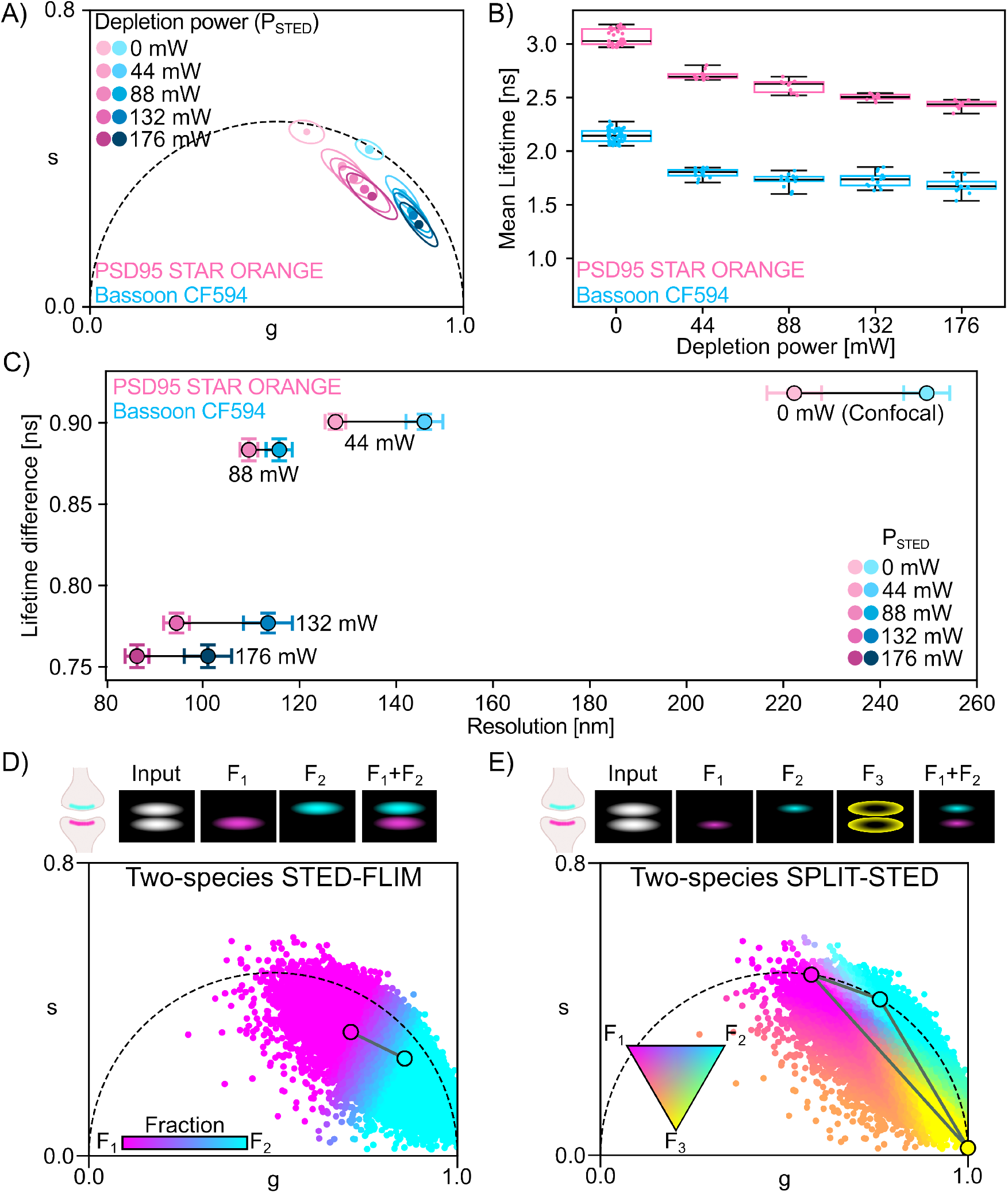
Effect of the depletion process on the fluorescence lifetime in two-species STED-FLIM. (A) Mean centroid and ellipse position of the phasor distributions for Bassoon-CF594 (Blue, N = 15 - 44 mW; 14 - 88 mW; 14 - 132 mW and 13 - 176 mW images) and PSD95-STAR ORANGE (Pink, 10 images per depletion laser power). (B) Mean lifetimes for each image plotted as median (horizontal black line) with boxes showing the quartiles and the whiskers showing the minima and maxima. The mean lifetime is reduced by the depletion laser power. (C) Relationship between the STAR ORANGE and CF594 lifetime difference and STED resolution. Shown is the mean lifetime difference and spatial resolutions for image pairs (N= 150- 44 mW; 140- 88 mW; 140- 132 mW and 130 - 176 mW image pairs of PSD95-STAR ORANGE and Bassoon-CF594) with the whiskers corresponding to the Standard Error of the Mean (SEM). (D) and (E) Top) Schematic of the components in the image space and Bottom) color-coded phasor plot based on fraction values for (D) two-species STED-FLIM and (E) two-species SPLIT-STED.

We propose to apply the SPLIT-STED strategy to two-species STED-FLIM. In two-species SPLIT-STED, we define three fractions, one for each fluorophore (*F*_1_, *F*_2_) and a third one for the short lifetime component associated with undepleted photons in the periphery of the super-resolved structures (*F*_3_)(Figure 2E). We exploit a strategy developed by Vallmitjana *et al*. [53] to separate *F*_3_ from *F* 1 and *F*_2_ using a linear system of equations (Equation 6). As in single-species SPLIT-STED, we obtain a resolution enhancement for both fluorophores. We define a reference triangle in phasor space joining the measured confocal lifetimes of both fluorophores to the point corresponding to both fluorophores’ short lifetime (Figure 2E, Figure S7). More precisely, in phasor space, the position of each pixel in a STED-FLIM image is analyzed in relation to the reference triangle to obtain a triplet of fractional components (*f*_1_,*f*_2_,*f*_3_) (Equation 6) that is then used to distribute its photons into three fractions (*F*_1_,*F*_2_,*F*_3_) (Figure 2E).

We first validated our two-species SPLIT-STED approach using a synthetic dataset created by combining STED-FLIM images of individually labeled proteins. These single-fluorophore images were acquired independently to serve as ground truth for performance evaluation[57](Methods, Figure 3A,B). As a baseline comparison, we used the two-species STED-FLIM phasor analysis approach to unmix the combined synthetic images (Figure 3C). We noticed that for increasing depletion powers, the borders of the longer lifetime fluorophore (*F*_1_) tend to be associated with the short lifetime fluorophore (*F*_2_) (Figure 3C, arrowheads), resulting in increased unmixing errors as measured with the NanoJ-SQUIRREL metric [54] (Figure 3E and Figure S8). Using the two-species SPLIT-STED approach, this border is assigned to the third fraction (*F*_3_), resulting in reduced unmixing errors for both fluorophores (Figure 3E-F and Supplementary Table S3). As for single-species SPLIT-STED, we show that the two-species SPLIT-STED strategy improves the spatial resolution for both structures (Figure 3F, Figure S9 and Table S4). Therefore, using synthetic dual-staining images, we confirmed that two-species SPLIT-STED results in improved spatial resolution and discrimination accuracy when unmixing the contribution of two fluorophores in STED-FLIM images.

**Figure 3.**
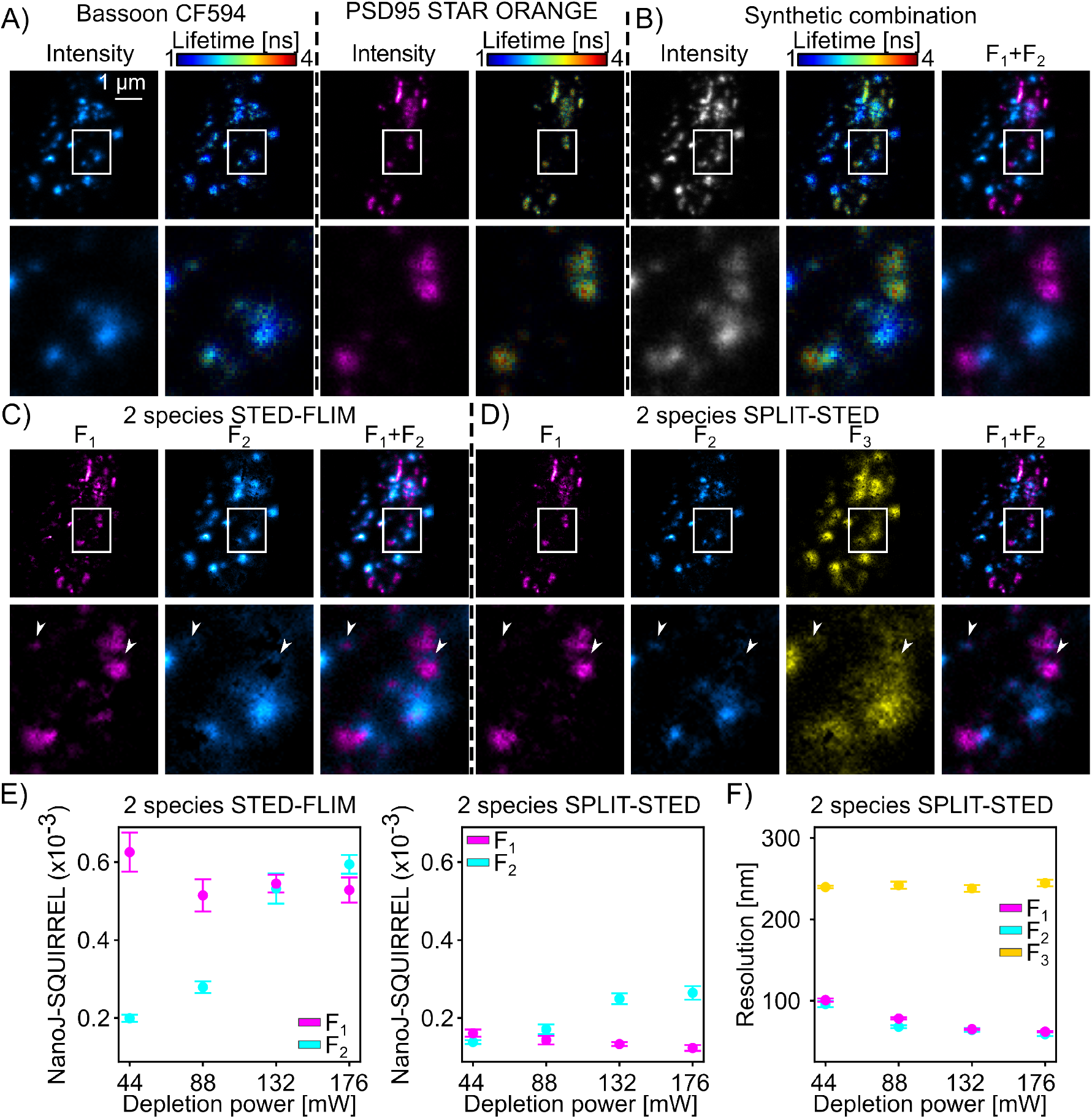
Validation of two-species SPLIT-STED on a synthetic dataset. (A) STED images of the synaptic proteins Bassoon labelled with CF594 and PSD95 labelled with STAR ORANGE. The images are color-coded for the photon counts (Bassoon-cyan, PSD95-magenta, left) and the measured fluorescence lifetime (1-4 ns, right). Inset (bottom) corresponds to the region identified by the white box. (B) Single-species STED-FLIM images are combined to generate a synthetic two-species STED-FLIM dataset with associated ground truth images for each fluorophore. (C) Phasor analysis of the STED-FLIM images with two fractions leads to unmixing errors at the border of the nanostructures labelled with STAR ORANGE (long lifetime). (D) In two-species SPLIT-STED the short lifetime border of STAR-ORANGE (arrowheads) is assigned to the third fraction (*F*_3_) improving unmixing accuracy. (E) Unmixing artifacts measured with NanoJ-SQUIRREL are larger for conventional two-species STED-FLIM phasor analysis (left) compared to two-species SPLIT-STED (right) (*F*_1_ - magenta, *F*_2_ - cyan, all p-values are shown in Table S3). (F) Two-species SPLIT-STED removes a low resolution component (*F*_3_) resulting in an improved resolution for both fluorophores (*F*_1_ and *F*_2_) in comparison to conventional STED-FLIM analysis (Figure S9 and all p-values are shown in Table S4). (E) and (F) The mean (points) and SEM (whiskers) are measured on the synthetic image dataset (n= 150- 44 mW; 140- 88 mW; 140- 132 mW and 130 - 176 mW image pairs).

We next applied two-species SPLIT-STED on real STED-FLIM images of pairs of neuronal proteins tagged with red or far-red fluorophores (Material and Methods). We considered pairs of STED-compatible fluorophores with non-overlapping confocal phasor distributions. Various protein combinations, including cytoskeletal proteins (βII-Spectrin and αTubulin) and synaptic proteins (PSD95, Homer1, Bassoon) were labelled and imaged in STED-FLIM mode within a single imaging channel (Figure 4, Figure 5 and Figure S10). The proteins of interest were labelled using primary and secondary antibodies. The fluorescence lifetime of the fluorophores were measured for each labelled protein as it is known to vary depending on the cellular environment [7], [9], [11]. We built a reference unmixing triangle in phasor space for each pair using single-species reference STED-FLIM images. Based on our results on single-species STED-FLIM and on the synthetic dataset, we adapted the depletion power to optimize the trade-off between spatial resolution and lifetime unmixing. On neuronal cultures stained for βII -Spectrin (STAR ORANGE) and Bassoon (CF594) we observed that two-species SPLIT-STED results in an improved spatial resolution of the membrane associated periodical lattice and pre-synaptic nanoclusters (Figure 4). By filtering the short-lifetime component, two-species SPLIT-STED minimizes the misassignment of photons from the periphery of the bnanostructures to the fluorophore with the shorter lifetime. In particular, for βII -Spectrin, shorter lifetime photons between the periodical structures are associated with fraction *F*_3_ by two-species SPLIT-STED, while they are assigned to the Bassoon *F*_2_ fraction when using conventional STED-FLIM phasor analysis. The improved resolution of two-species SPLIT-STED is also necessary to resolve the filaments of αTubulin stained with Alexa Fluor 647 and Bassoon stained with STAR 635P (Figure 5). Alexa Fluor 647 is associated with high photobleaching rates in STED microscopy which requires the use of lower depletion powers in comparison to other conventional far-red STED-compatible fluorophores. It limits its application for two-color STED microscopy. Here, we show that using two-species SPLIT-STED allows the reduction of the required depletion power (44 mW), while maintaining sufficient spatial resolution to resolve αTubulin filaments and Bassoon nanoclusters (Figure 5). Since the third fraction is discarded from the unmixed image, a sharper separation of the Tubulin filaments and of the Bassoon nanoclusters can be noted on intensity profiles (Figure 5E-F). We observed a similar behaviour for βII Spectrin and Bassoon labelled with Alexa Fluor 647 and STAR 635P respectively (Figure S10C). We next tested the applicability of two-species SPLIT-STED on pairs of synaptic proteins (Bassoon-PSD95 and Bassoon-Homer1)(Figure S10A-B), showing similar intensity distributions and often overlapping signals due to the proximity (*∼*100 nm) of the protein nanoclusters [37]. In accordance with the results obtained on the synthetic dataset, two-species SPLIT-STED improves the spatial resolution and unmixing performance in comparison to conventional STED-FLIM phasor analysis for red and far-red fluorophores (Figure S10). Thus, we demonstrate that two-species SPLIT-STED yields precise separation two fluorophores in real STED-FLIM images. It can be used on pairs of spectrally overlapping fluorophores and reduces the required laser power in comparison to conventional STED-FLIM imaging.

**Figure 4.**
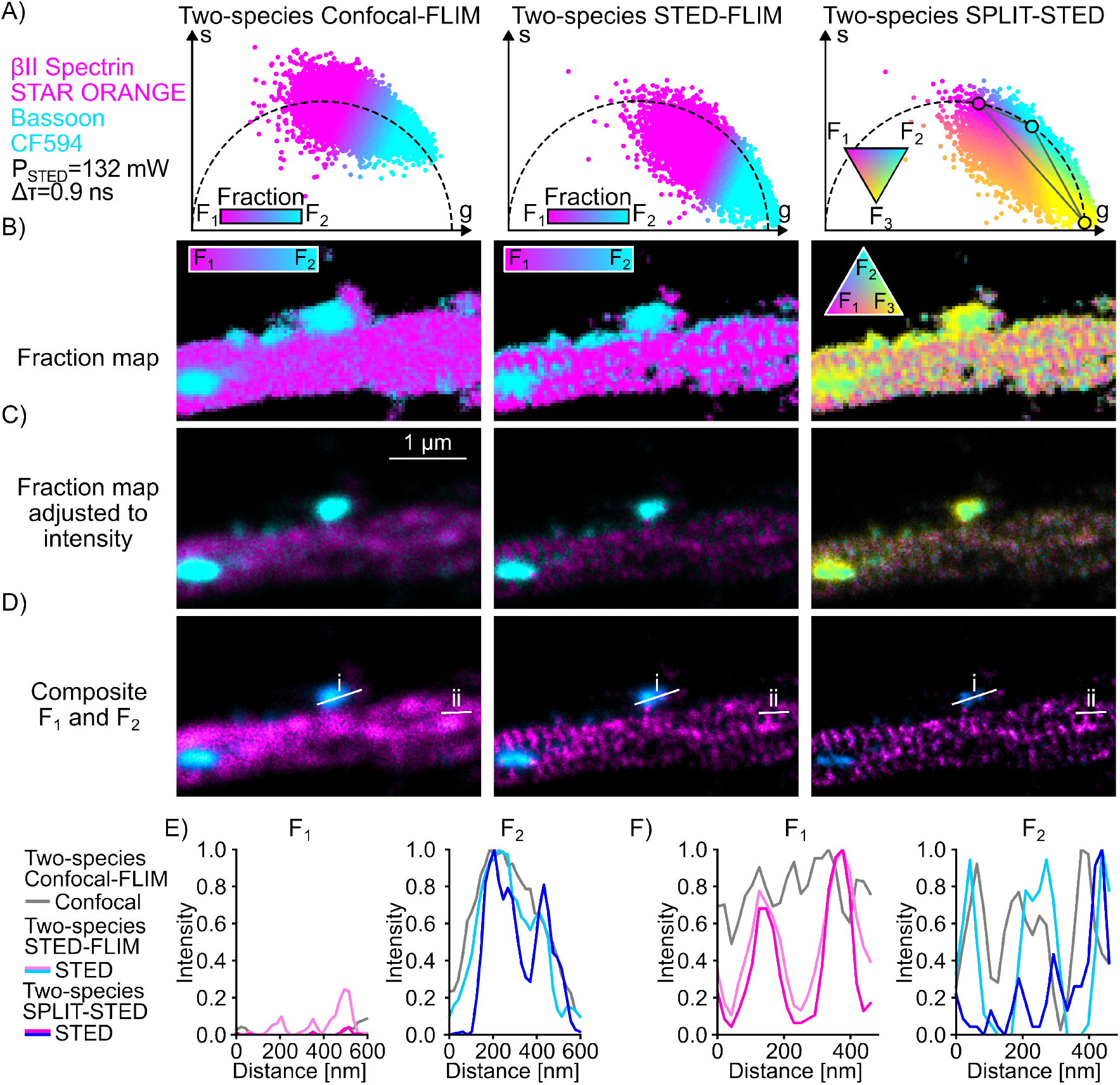
Two-species SPLIT-STED of neuronal nanostructures labelleled with red fluorophores. Representative unmixed image using the two-species STED-FLIM and two-species SPLIT-STED approaches of Bassoon CF594 and βII-Spectrin STAR ORANGE (P_STED_ = 132 mW). (A) Phasor plots of the confocal (left) and STED (middle and right) images color-coded based on the assigned fraction for each pixel (Methods). For each method, we show the pixel-wise fraction map (B), which is adjusted to correspond to the intensity distribution (C). The resulting unmixed images (D) are used to obtain the intensity profiles shown in (E) and (F). (E) Two-species SPLIT-STED resolves Bassoon nanoclusters that could not be distinguished in confocal or conventional STED-FLIM images (profile i in panel (D)). (F) βII-Spectrin rings are clearly resolved in *F*_1_ in both SPLIT-STED and STED-FLIM images but the space between the rings is assigned to *F*_2_ in STED-FLIM, while being filtered out in SPLIT-STED (profile ii in panel (D).

**Figure 5.**
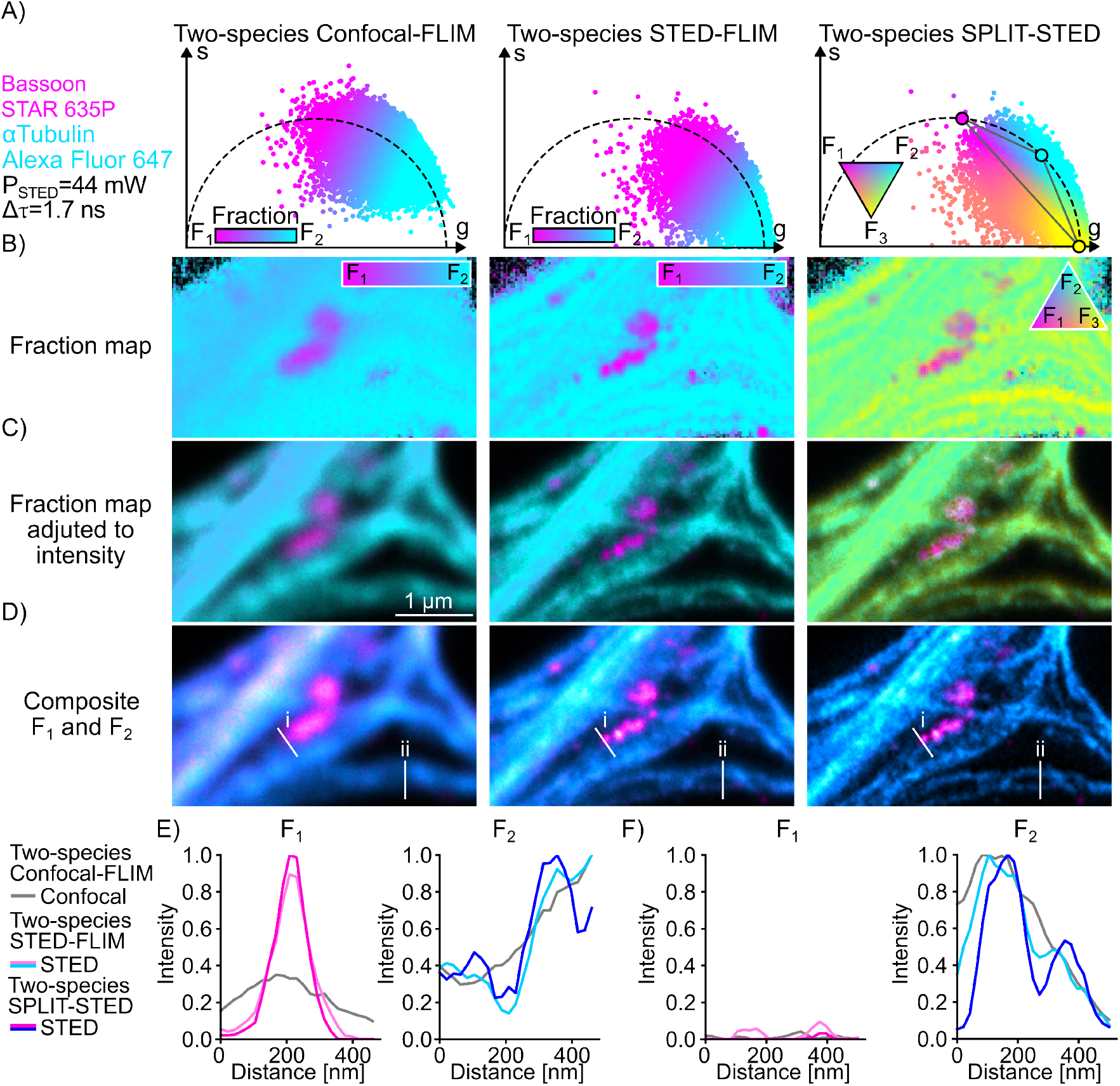
Two-species SPLIT-STED of neuronal nanostructures labelleled with far-red fluorophores. Representative unmixed image using the two-species STED-FLIM and two-species SPLIT-STED approaches of αTubulin Alexa Fluor 647 and Bassoon STAR 635P imaged with P_STED_ = 44 mW. (A) Phasor plots of the confocal (left) and STED (middle and right) images color-coded based on the assigned fraction for each pixel (Methods). For each method we show the pixel-wise fraction map (B), which is adjusted to correspond to the intensity distribution (C). The resulting unmixed images (D) are used to obtain the intensity profiles shown in (E) and (F). (E) Two-species SPLIT-STED provides a sharper discrimination of two αTubulin filaments surrounding a Bassoon cluster (profile i in panel (D)) (F) Two-species SPLIT-STED better resolves two neighbouring αTubulin filaments (profile ii in panel (D)).

## 4 Conclusion

We developed an analytical approach, two-species SPLIT-STED, for the unmixing of STED-FLIM images based on the law of linear combination in phasor space. Two-species SPLIT-STED associates photons from undepleted fluorophores at the border of the PSF to a third fraction. It thereby reduces the unmixing artifacts of two-species STED-FLIM, while increasing the measured spatial resolution. This analytical approach is shared as open-source code together with a dataset of STED-FLIM images of red and far-red dyes [57]. We validated the approach first on a synthetic dataset, which provided single-species ground truth images, allowing for a robust quantification of unmixing artifacts and spatial resolution. We also showed on various neuronal protein pairs labelled with red and far-red emitting fluorophores, that two-species

SPLIT-STED leads to improved unmixing results on real biological samples. The improved spatial resolution at lower depletion power provided by this approach could be further applied to live-cell STED imaging. It could also be integrated to multi-channel STED-FLIM imaging, given that appropriate fluorophore pairs are available [7], [13], [14], [30].

For a given targeted spatial resolution, two-species SPLIT-STED reduces the required depletion power in comparison to two-species phasor-FLIM analysis. This allows a better lifetime separation to be maintained between the chosen fluorophores, which is known to be diminished by high depletion powers [14].

Nevertheless, a careful optimization of fluorophores and protein pairs, taking into account both the measured lifetime at the desired depletion power as well as the relative brightness of both fluorophores is required [7], [9], [11], [30], [34]. Since the lifetime of fluorophores varies depending on which protein it labels, this optimization needs to be repeated for each potential protein combination. With the constantly improving fluorescence labelling strategies and fluorescent probes, analytical approaches, such as two-species SPLIT-STED, could contribute to expanding multiplexed super-resolution microscopy in fixed and living samples.

## Acknowledgments

We thank the Neuronal Cultures Platform of the CERVO Brain Research Centre for the preparation of the primary neuronal cultures. We thank Andreas Schönle and Markus Köhler from Abberior Instruments GmbH for their assistance with the integration of the control of the Time Tagger in the Imspector Software. We thank Annette Schwerdtfeger for careful proofreading of the manuscript.

## Research funding

Funding was provided by grants from the Natural Sciences and Engineering Research Council of Canada (RGPIN-2017-06171, P.D.K., RGPIN-2019-06704 F.L.C.), Fonds de Recherche Nature et Technologies (FRQNT), Team Grant (2021-PR-284335 to F.L.C, C.G., and P.D.K.), Canadian Institutes of Health Research (CIHR) (202109PJT-471107-NSB-CFBA-12805 to F.L.C. and P.D.K.), Neuronex Initiative (National Science Foundation 2014862, Fond de recherche du Québec - Santé 295824 to F.L.C. and P.D.K.), the Canadian Foundation for Innovation (32786 to P.D.K. and 39088, F.L.C.), and Canada First Research Excellence Fund Sentinelle Nord. F.L.C. is a Canada Research Chair Tier II (CRC-2023-00385, F.L.C.) and C.G. is a CIFAR AI-Chair.

A.B. was supported by an excellence scholarship from NSERC. A.B. and A.M.G. were awarded excellence scholarships from the FRQNT strategic cluster UNIQUE. M.L. was supported by an excellence scholarship from the Institute for Intelligence and Data.

## Author Contributions

A.D, A.M.G., A.O., M.L implemented the MLE and phasor-based analysis approaches for Confocal-FLIM and STED-FLIM microsocopy. A.D., M.L., and A.O.. performed the comparative lifetime fitting analysis. A.D. developed the two-species SPLIT-STED analysis approach. A.B. implemented image visualisation approaches. A.B. and A.D. implemented the performance evaluation approaches. A.D. generated the synthetic image dataset. A.D. and A.B developed the automated STED-FLIM acquisition pipeline. A.O., and A.B. integrated the Time Tagger to the STED microscope. A.D. and J.G.S.S. prepared the samples.

A.D. acquired the images on the STED microscope. A.D. and F.L.C designed the experimental protocols.

A.D. and F.L.C. wrote the manuscript. P.D.K, C.G and F.L.C co-supervised the project. All authors have accepted responsibility for the entire content of this manuscript and approved its submission.

## Data Availability

The Confocal- and STED-FLIM Images of Neuronal Proteins dataset is available at https://zenodo.org/records/15438495[57].

## Software Availability

The code used to generate the results and figures is available in a public Github repository at https://github.com/FLClab/2-Species-SPLIT-STED

## List of Acronyms

STED: STimulated Emission Depletion
FLIM: Fluorescence Lifetime Imaging Microscopy
SPLIT: Separation of Photons by LIfetime Tuning
PSF: Point Spread Function
TCSPC: Time Correlated Single Photon Counting
DIV: days in vitro
PAB: primary antibodies
SAB: secondary antibodies
PFA: paraformaldehyde
PBS: Phosphate-Buffered Saline
NGS: normal goat serum
RT: room temperature
APD: Avalanche Photodiode Detectors
IRF: Instrument Response function
FWHM: Full Width at Half Maximum
MLE: Maximum Likelihood Estimation
ROI: regions of interest
SLSQP: Sequential Least Squares Programming
CWF: Complex Wavelet Filter
SQUIRREL: Super-resolution QUantitative Image Rating and Reporting of Error Locations
SE: Squared Error
SEM: Standard Error of the Mean
SNR: signal to noise ratio
STD: standard Deviation
BFP: Back Focal Plane
CW: Continuous Wave
FCS: Fluorescence Correlation Spectroscopy

## Notes

https://doi.org/10.5281/zenodo.15438495

https://github.com/FLClab/2-Species-SPLIT-STED

### Competing Interest Statement

The authors have declared no competing interest.

