## Supplementary Material for "Enhancing STED Microscopy via Fluorescence Lifetime Unmixing and Filtering in Two-Species SPLIT-STED"

| Primary Antibodies | | | | |
|---|---|---|---|---|
| **Antibody** | **Supplier** | **Catalogue no.** | **Dilution** | **Ref.** |
| Mouse anti-PSD95 | Abcam | MA1-045 | 1 : 500 | [37], [58] |
| Mouse anti-Bassoon | Enzo | ADI-VAM-PS003 | 1 : 500 | [37], [59], [60] |
| Rabbit anti-Bassoon | Synaptic Systems | 141003 | 1 : 500 | [37], [59] |
| Rabbit anti-Homer1 | Synaptic Systems | 160003 | 1 : 500 | [61] |
| Mouse anti-β2-Spectrin | BD Biosciences | 612563 | 1 : 250 | [39], [62] |
| Mouse anti-α-Tubulin | Sigma-Aldrich | T5168 | 1 : 500 | [63] |

| Secondary Antibodies | | | |
|---|---|---|---|
| **Antibody** | **Supplier** | **Catalogue no.** | **Dilution** |
| Goat anti-Rabbit CF594 | Sigma-Aldrich | SAB4600407 | 1 : 500 |
| Goat anti-Mouse CF594 | Sigma-Aldrich | SAB4600321 | 1 : 500 |
| Goat anti-Rabbit STAR ORANGE | Abberior | STORANGE-1002 | 1 : 250 |
| Goat anti-Mouse STAR ORANGE | Abberior | STORANGE-1001 | 1 : 250 |
| Goat anti-Rabbit STAR 635P | Abberior | ST635P-1001 | 1 : 250 |
| Goat anti-Mouse Alexa Fluor 647 | Jackson Immunoresearch | 115-605-166 | 1 : 250 |

**Table S1.** Antibodies used for immunostaining

| Red channel | | |
|---|---|---|
| **Parameter** | **Confocal-FLIM** | **STED-FLIM** |
| **Excitation wavelength** | 561 nm | 561 nm |
| **Excitation power** | 2.6 µW | 2.6 µW |
| **Depletion wavelength** | | 775 nm |
| **Depletion powers** | | 44, 88, 132, 176 mW |
| **Detection wavelengths** | 605-625 nm | 605-625 nm |
| **Detection time bins** | 250 bins of 80 ps | 250 bins of 80 ps |
| **Pixel dwelltime** | 15 µs | 15 µs |
| **Line steps** | 6 | 12 or 25 (Fig 3) |
| **Pixel size** | 20 nm | 20 nm |
| **Pinhole size** | 1.0 A.U. | 1.0 A.U. |

| Far-red channel | | |
|---|---|---|
| **Parameter** | **Confocal-FLIM** | **STED-FLIM** |
| **Excitation wavelength** | 640 nm | 640 nm |
| **Excitation power** | 1 µW | 2 µW |
| **Depletion wavelength** | | 775 nm |
| **Depletion powers** | | 22,44,66,88 mW |
| **Detection wavelengths** | 650-720 nm | 655-720 nm |
| **Detection time bins** | 250 bins of 80 ps | 250 bins of 80 ps |
| **Pixel dwelltime** | 15 µs | 15 µs |
| **Line steps** | 8 | 10 |
| **Pixel size** | 20 nm | 20 nm |
| **Pinhole size** | 1.0 A.U. | 1.0 A.U. |

**Table S2.** Imaging parameters for acquisition of FLIM images

| Depletion Power | Posthoc Dunn's test (p-values)<br>Two-species STED-FLIM vs Two-species SPLIT-STED<br>F1 | F2 |
|---|---|---|
| 44 mW | $\mathbf{2.04 \times 10^{-22}}$ | $\mathbf{9.14 \times 10^{-8}}$ |
| 88 mW | $\mathbf{1.47 \times 10^{-19}}$ | $\mathbf{5.21 \times 10^{-14}}$ |
| 132 mW | $\mathbf{1.343 \times 10^{-40}}$ | $\mathbf{1.04 \times 10^{-16}}$ |
| 176 mW | $\mathbf{1.98 \times 10^{-29}}$ | $\mathbf{1.22 \times 10^{-28}}$ |

**Table S3.** Comparison of NanoJ-SQUIRREL error values for two-species STED-FLIM and two-species SPLIT-STED on the synthetic dataset for different depletion powers. The p-values are obtained by a posthoc Dunn's test following a Kruskal–Wallis H test to compare the distributions of each method at each depletion power (Methods). The main results are reported in Figure 3E. Bold: significant difference between groups (p <0.05)

Posthoc Dunn's test (p-values)

| Depletion Power | Input vs Two-species STED-FLIM<br>$F_1$ | $F_2$ | Input vs Two-species SPLIT-STED<br>$F_1$ | $F_2$ |
|---|---|---|---|---|
| 44 mW | $\mathbf{1.45 \times 10^{-16}}$ | $\mathbf{2.32 \times 10^{-18}}$ | $\mathbf{1.58 \times 10^{-28}}$ | $\mathbf{3.38 \times 10^{-11}}$ |
| 88 mW | $\mathbf{1.56 \times 10^{-37}}$ | $\mathbf{1.98 \times 10^{-9}}$ | $\mathbf{5.07 \times 10^{-39}}$ | $\mathbf{6.42 \times 10^{-26}}$ |
| 132 mW | $\mathbf{1.39 \times 10^{-22}}$ | $3.53 \times 10^{-1}$ | $\mathbf{3.43 \times 10^{-50}}$ | $\mathbf{1.21 \times 10^{-33}}$ |
| 176 mW | $\mathbf{3.23 \times 10^{-26}}$ | $1.26 \times 10^{-1}$ | $\mathbf{4.29 \times 10^{-38}}$ | $\mathbf{5.07 \times 10^{-52}}$ |

Posthoc Dunn's test (p-values)

| Depletion Power | Two-species STED-FLIM vs Two-species SPLIT-STED<br>$F_1$ | $F_2$ |
|---|---|---|
| 44 mW | $\mathbf{4.82 \times 10^{-3}}$ | $\mathbf{2.63 \times 10^{-53}}$ |
| 88 mW | $7.92 \times 10^{-1}$ | $\mathbf{2.33 \times 10^{-61}}$ |
| 132 mW | $\mathbf{3.07 \times 10^{-7}}$ | $\mathbf{9.70 \times 10^{-39}}$ |
| 176 mW | $\mathbf{4.56 \times 10^{-6}}$ | $\mathbf{5.77 \times 10^{-30}}$ |

**Table S4.** Comparison of spatial resolutions obtained on the synthetic dataset with two-species STED-FLIM and two-species SPLIT-STED for different depletion powers. The p-values are obtained by a posthoc Dunn's test following a Kruskal–Wallis H test to compare the distributions of each method at each depletion power (Methods). The main results are reported in Figures 3F and S9. Bold: significant difference between groups (p <0.05)

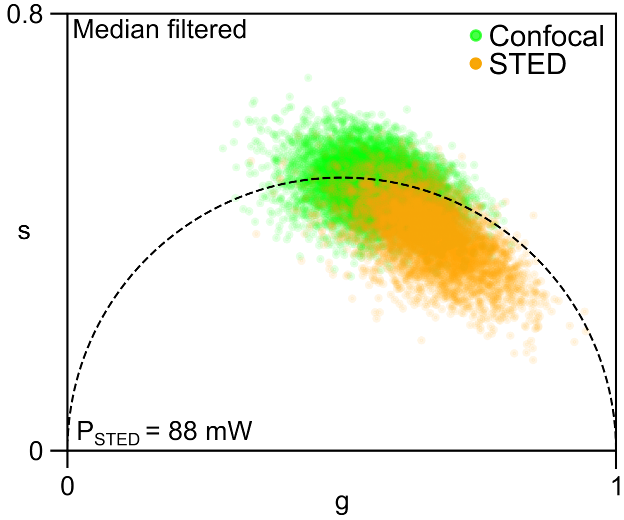


**Fig. S1. Phasor distribution of a Confocal-(green) and STED-(orange) FLIM image of Homer1 STAR ORANGE.**

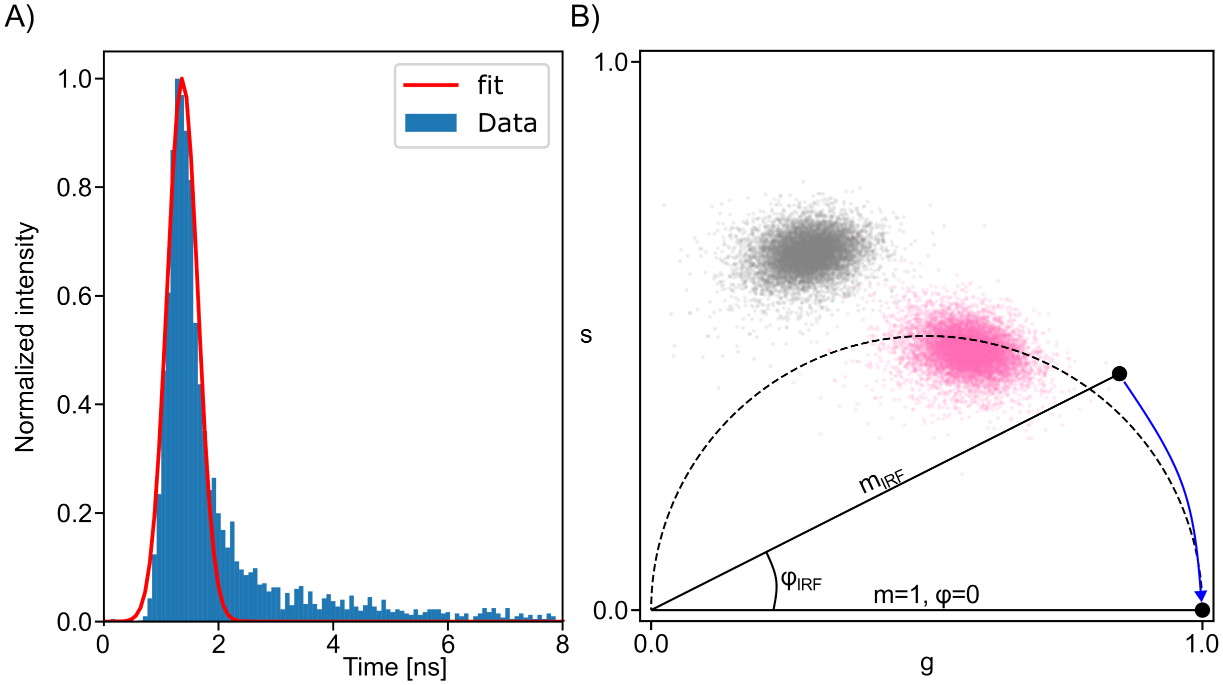


**Fig. S2. Calibration of phasor plots using IRF measurement**
(A) Histogram of the IRF measurement (blue) and gaussian fit (red). (B) Phasor distribution of a Confocal-FLIM image of PSD95 STAR ORANGE before (grey) and after (pink) calibration based on a measurement of the IRF.

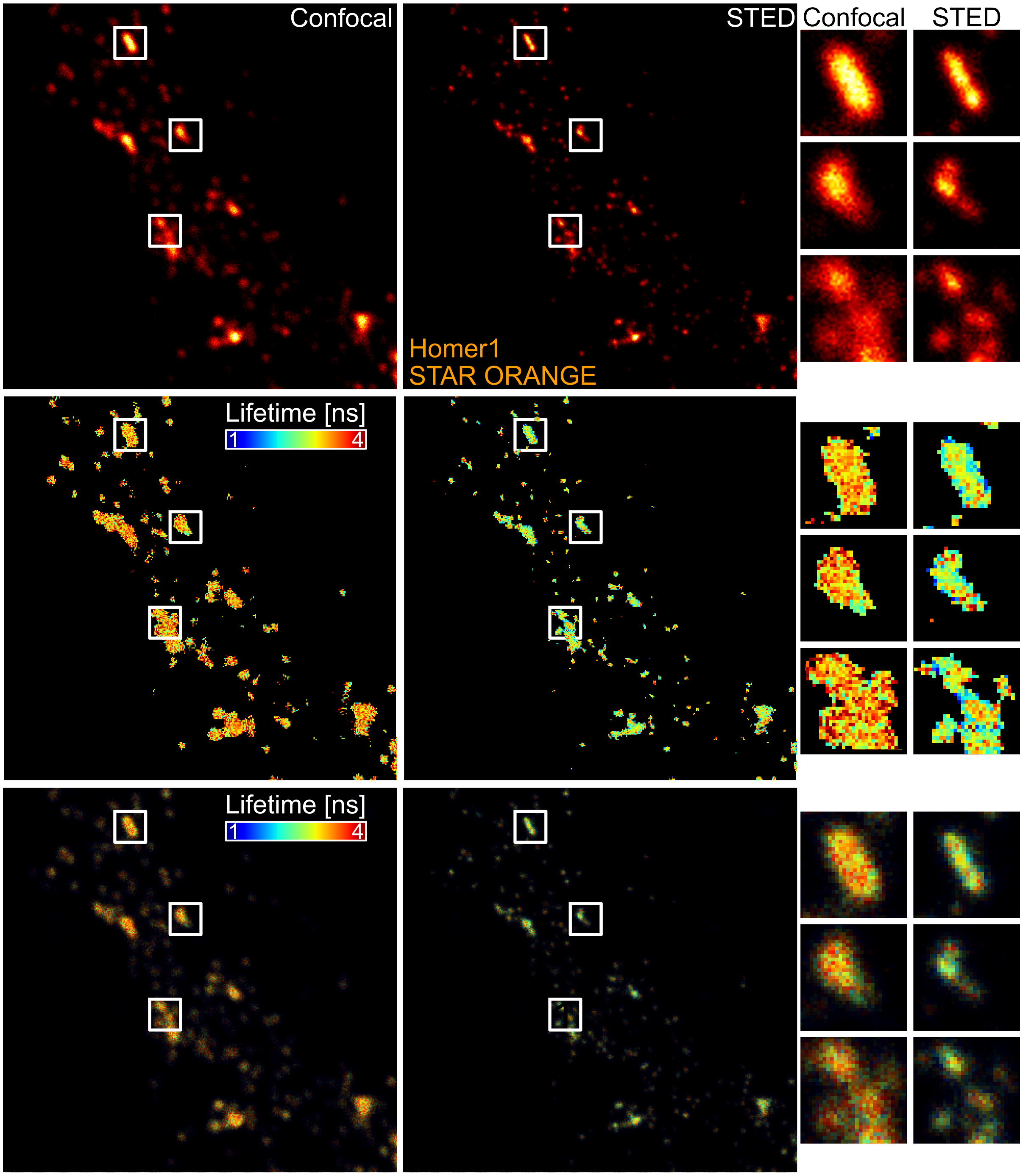


**Fig. S3. Spatial distribution of lifetime in confocal and STED images**
Confocal and STED images of the pre-synaptic protein Homer1 labelled with STAR ORANGE color-coded for Top) Pixel intensity. Middle) mean lifetime obtained with mono-exponential histogram fitting. Bottom) intensity image color-coded for the lifetime distribution.

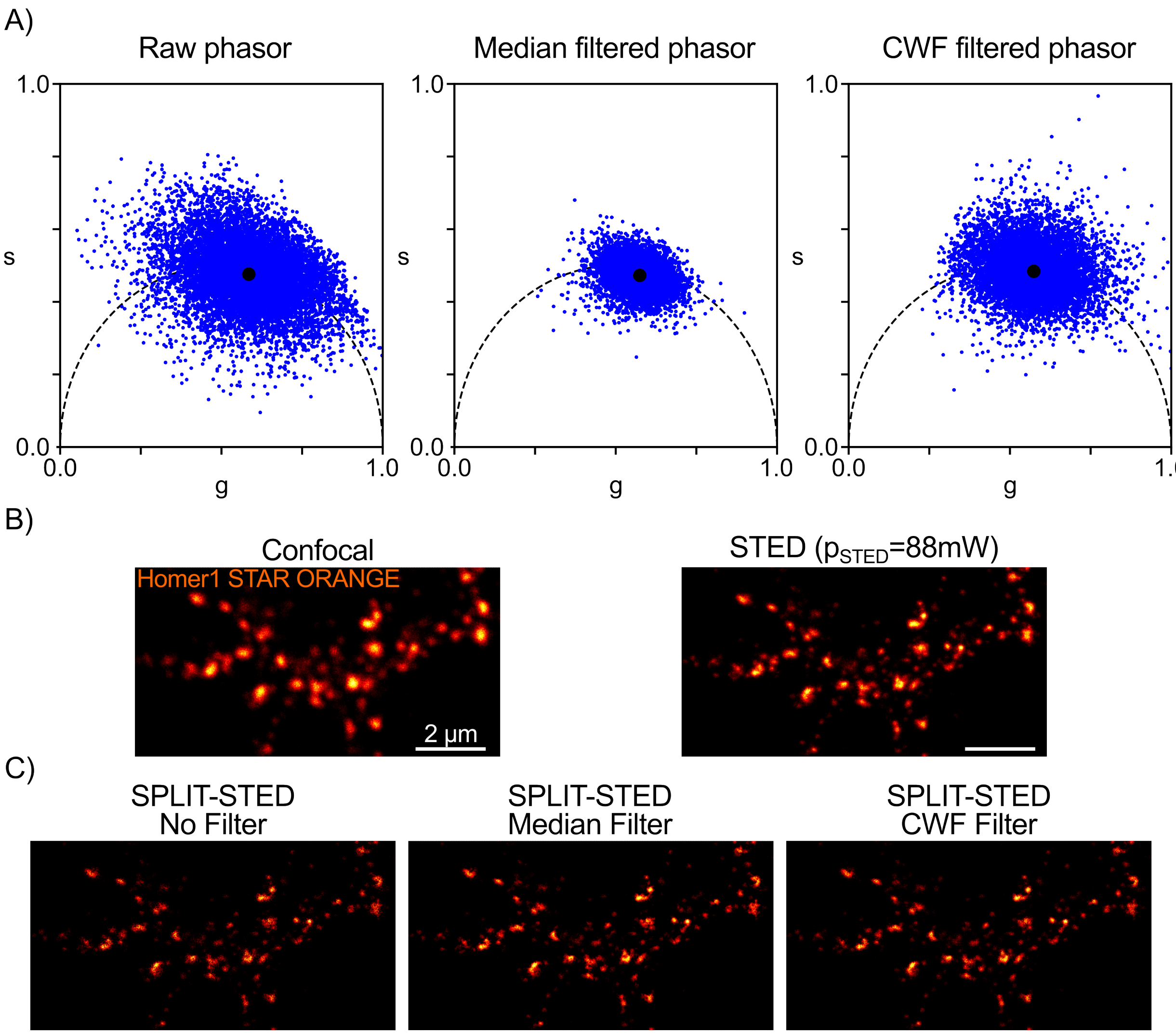


**Fig. S4. Filtering methods in phasor space**
(A) Raw phasor distribution of a Confocal-FLIM image of Homer1 STAR ORANGE before filtering (left), with Median filtering (middle) and with CWF filtering (right). (B) Input intensity images of Homer1 STAR ORANGE confocal (left) and STED (right). (C) Comparison of SPLIT-STED images for the three filtering approaches shown in (A).

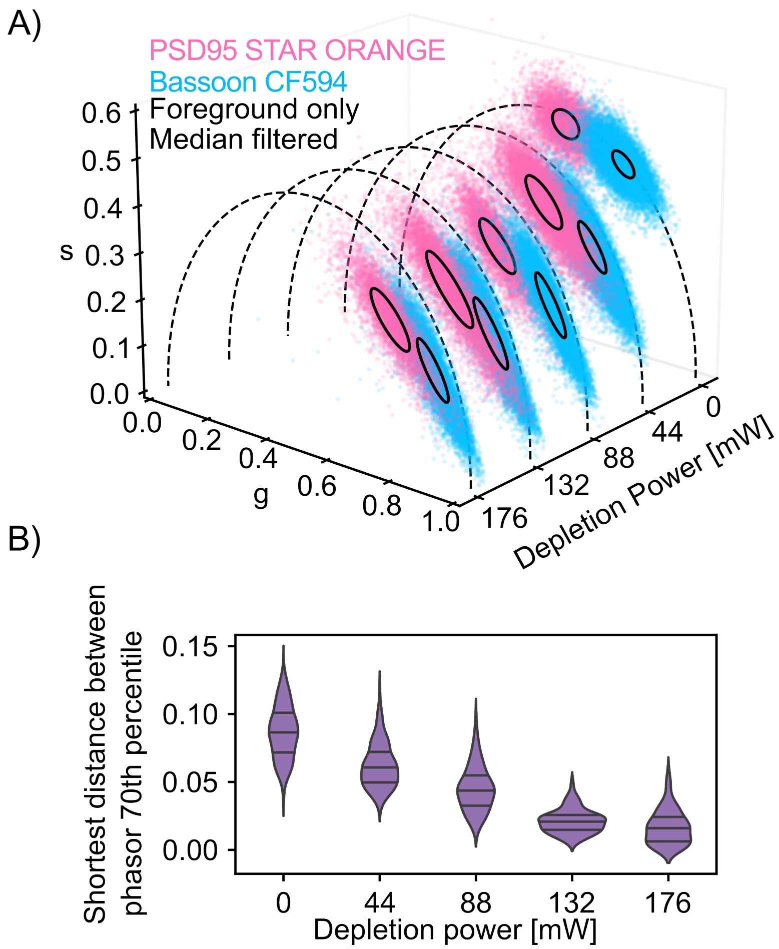


**Fig. S5. Phasor distributions of STED-FLIM images acquired with varying depletion powers** (A) Representative phasor distributions from single-species images acquired with varying depletion powers. When plotted in the same phasor space, the fluorescence lifetime of Bassoon-CF594 (blue) and PSD95-STAR ORANGE (pink) show increased ellipticity and overlap for increasing STED depletion power. Ellipses (black) represent the 70th percentile of the covariance distribution. (B) Violin plot of the shortest distance between the borders of 70th percentile ellipses. The distance was calculated for each synthetic pair of single-species STED-FLIM images from PSD95-STAR ORANGE and Bassoon CF594. The STED depletion laser significantly affects the shortest distance between the lifetime distributions for all depletion powers when compared to the confocal-FLIM lifetime distributions (p-value from Posthoc Dunn <0.001, ***). Horizontal lines are quartiles.

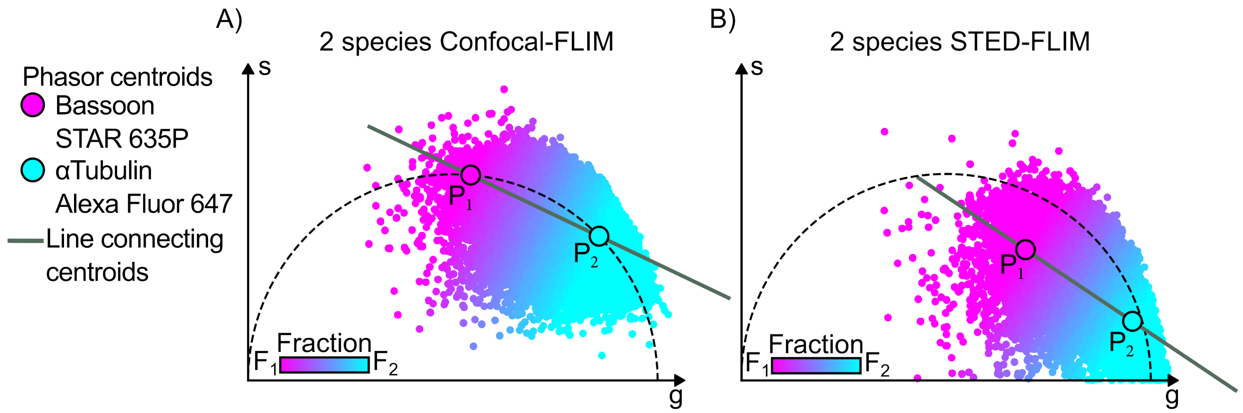


**Fig. S6. Generation of the reference lines for two-species Confocal- and STED-FLIM**
The centroids of the lifetime distributions in phasor space from single species images of Bassoon STAR 635P (pink) and αTubulin Alexa Fluor 647 (blue) are used to define the pure species reference points $P_1$ and $P_2$ on the universal semicircle. The reference line for (A) two-species Confocal-FLIM and (B) two-species STED-FLIM ($P_{STED}$ = 44 mW) is built using the $P_1$ and $P_2$ reference points. Each point of the phasor distribution of the two-species image is projected onto the reference line. A fraction value is assigned to each point based on its position on the line (color-code).

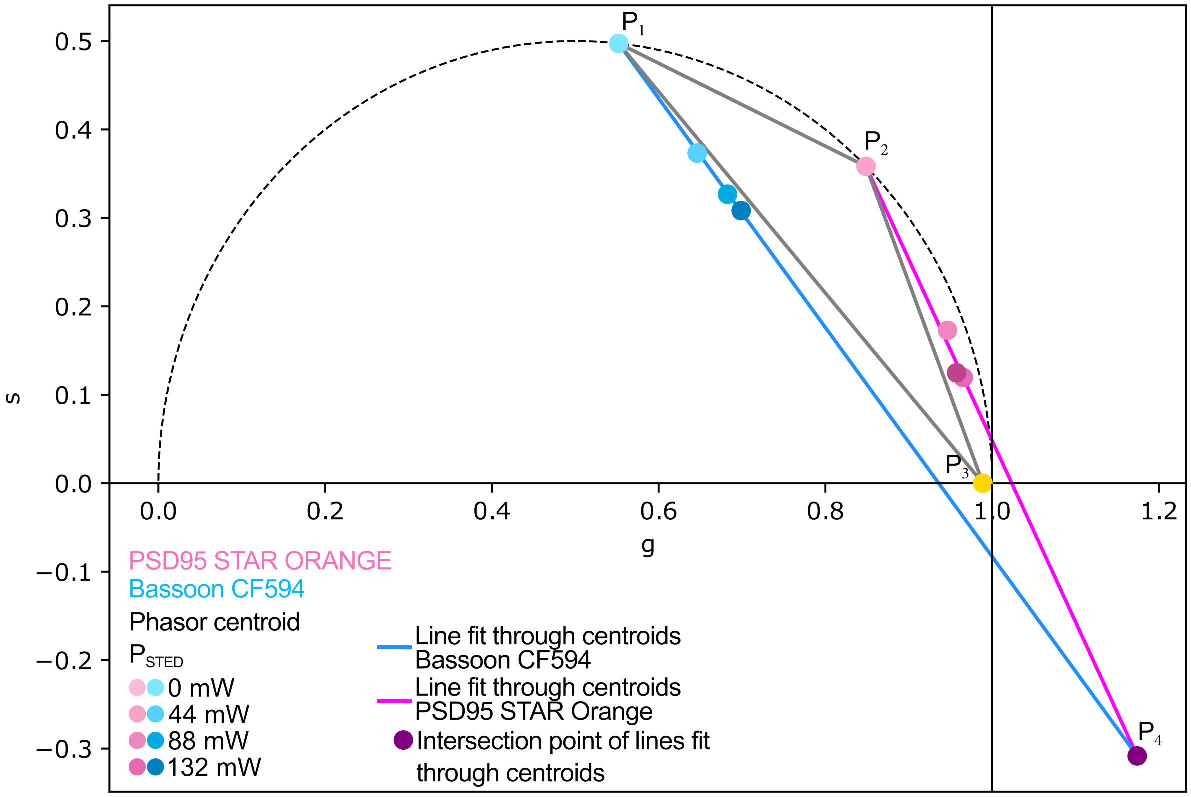


**Fig. S7. Generation of a reference triangle for two-species SPLIT-STED**
The centroids of the lifetime distributions in phasor space from single species confocal images of PSD95-STAR ORANGE (pink) and Bassoon CF594 (blue) are used to define the pure species reference points $P_1$ and $P_2$ on the universal semicircle. A linear fit on the centroid of the lifetime distributions from STED-FLIM images acquired at different depletion powers is used to define an intersection between both trajectories ($P_4$). The reference point ($P_3$) for the third fraction $F_3$ is defined as the nearest point to $P_4$ within the universal semicircle. The reference triangle for two-species SPLIT-STED is built using the $P_1$, $P_2$ and $P_3$ reference points.

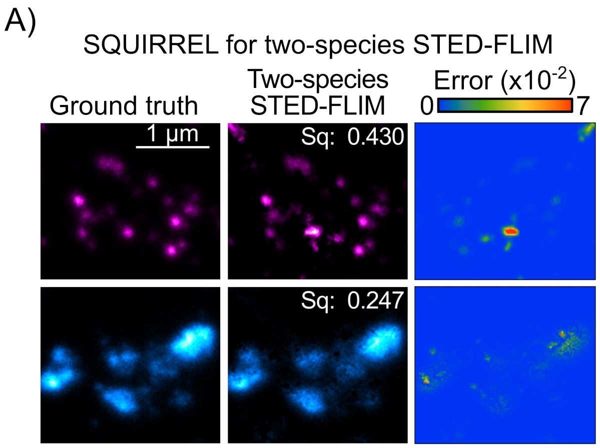


**Fig. S8. Maps of NanoJ-SQUIRREL error score**

Representative maps of NanoJ-SQUIRREL error score for $P_{STED} = 88\,\mathrm{mW}$ obtained on pairs of PSD95 STAR ORANGE (magenta) and Bassoon CF594 (cyan) images from the synthetic dataset for two-species (A) STED-FLIM and (B) SPLIT-STED. Ground truth image (left), unmixed image (middle) and error map (right).

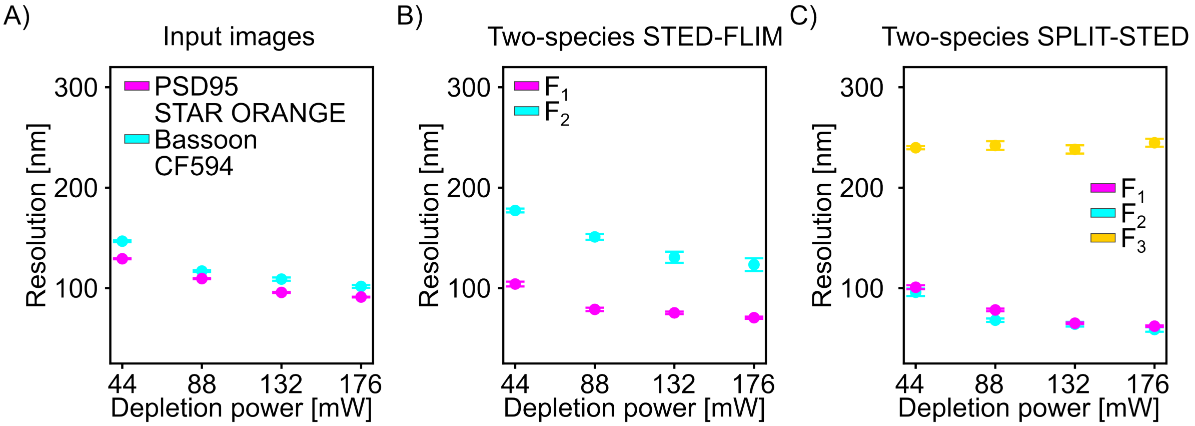


**Fig. S9. Spatial resolution for different depletion power for two-species STED-FLIM and SPLIT-STED**. The spatial resolution is measured on the synthetic dataset using a decorrelation approach for the synthetic ground truth (A), the unmixed STED-FLIM (B) and SPLIT-STED images (C). SPLIT-STED improves the spatial resolution for the fraction $F_1$ and $F_2$ and filters out the low resolution $F_3$ component. Synthetic images were created for pairs of Bassoon-CF594 (56 images, 15- 44 mW; 14- 88 mW;14- 132 mW and 13 - 176 mW) and PSD95-STAR ORANGE (40 images, 10 images per depletion power). Shown is the mean and SEM for each depletion power. Panel (C) is the same as Figure 3F).

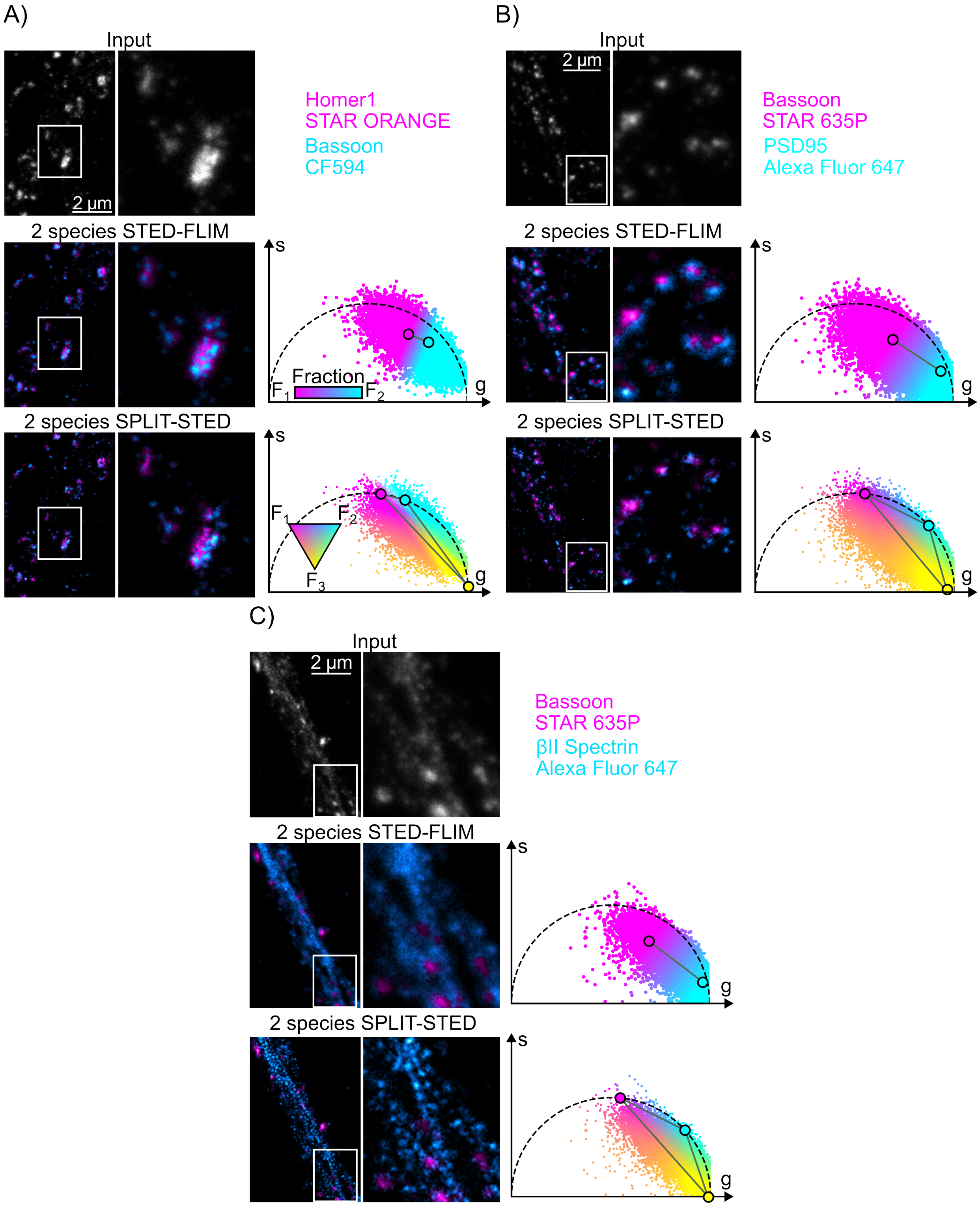


**Fig. S10. Unmixing of different neuronal protein pairs in real images**
Two-species SPLIT-STED is applied to different pairs of neuronal proteins labelled with red and far-red emitting fluorophores. (A) Bassoon CF594 Homer1 STAR ORANGE imaged with $P_{STED} = 132\,\text{mW}$ (B) PSD95 Alexa Fluor 647 and Bassoon STAR 635P imaged with $P_{STED} = 44$mW and (C) βII-Spectrin Alexa Fluor 647 and Bassoon STAR 635P imaged with $P_{STED} = 44$mW. Intensity image (top) and unmixed images with two-species STED-FLIM (middle) and two-species SPLIT-STED approaches (bottom). Phasor plots color-coded by the assigned fraction at each pixel are shown for both methods.
